# Prodrug suicide gene therapy for cancer targeted intracellular by mesenchymal stem cell exosomes

**DOI:** 10.1101/281808

**Authors:** Ursula Altanerova, Jana Jakubechova, Katarina Benejova, Petra Priscakova, Martin Pesta, Pavel Pitule, Ondrej Topolcan, Juraj Kausitz, Martina Zdurjencikova, Vanda Repiska, Cestmir Altaner

## Abstract

Human tumor trophic mesenchymal stem cells (MSCs) isolated from various tissues and MSCs engineered to express the yeast cytosine deaminase::uracil phosphoribosyl transferase suicide fusion gene (yCD::UPRT‒MSCs) released exosomes in conditional medium (CM). Exosomes from all tissue specific yCD::UPRT‒MSCs contained mRNA of the suicide gene in the exosome’s cargo. When the CM was applied to tumor cells, the exosomes were internalized by recipient tumor cells and in the presence of the prodrug 5‒fluorocytosine (5‒FC) effectively triggered dose‒dependent tumor cell death by endocytosed exosomes via an intracellular conversion of the prodrug 5‒FC to 5‒fluorouracil. Exosomes were found to be responsible for the tumor inhibitory activity. MSCs transduced with the Herpes simplex virus thymidine kinase gene released exosomes causing death of tumor cells in the presence of ganciclovir. The presence of microRNAs in exosomes produced from naive MSCs and corresponding transgene transduced MSCs did not differ significantly. microRNAs from yCD::UPRT‒MSCs were not associated with therapeutic effect. MSC suicide gene exosomes represent a new class of tumor cell targeting drug acting intracellular with curative potential.

## Introduction

Mesenchymal stem cells (MSCs) possess the ability to migrate to the site of injury and secrete a variety of factors capable of a number of functions inducing and supporting regenerative processes in the damaged tissue, inducing angiogenesis, protecting cells from apoptotic cell death and modulating the immune system (reviewed in Shi *et al*, 2017). It has been increasingly observed that the transplanted MSCs did not necessarily engraft and differentiate at the site of injury but might exert their therapeutic effects in a paracrine/endocrine fashion inducing an endogenous reparatory processes (Leibacher & Henschler, 2016). Rich secretome from MSCs and extracellulary released exosomes participate in the paracrine cross-talk between cells by delivering molecular information that leads to biological consequences. MSC-derived exosomes function as paracrine mediators in tissue repair and recapitulate to a large extent the therapeutic effects of parental MSCs (Phinney & Pittenger, 2017). The tumor, being a “wound that does not heal” (Dvorak, 1986]), attracts MSCs. The MSCs home in on the tumor and, together with other cells, form the tumor stroma. In vitro results suggested that the SDF-1-CXCR4 and HGF-c-met axes, along with MMPs, may be involved in recruitment of expanded MSCs to damaged tissues (<u>Son</u> *et al*, 2006). MSC tumor tropism inspired us to develop a prodrug gene therapy mediated by various types of MSCs. We have developed two prodrug suicide gene therapy systems: MSCs engineered to express fused yeast cytosine deaminase::uracil phosphoribosyl transferase (yCD::UPRT) with 5-fluorocytosine (5-FC) as a prodrug – yCD::UPRT-MSC/5-FC system) (Kucerova et al., 2007) and MSCs expressing thymidine kinase of the Herpes simplex virus with ganciclovir as a prodrug – tk*HSV*-MSC/ganciclovir system, (Matuskova *et al*, 2010). Both prodrug gene therapeutic systems express enzymes for the conversion of a non-toxic prodrug to an efficient cytotoxic compound from a cell DNA integrated provirus. MSCs transduced with suicide genes had retained their tumor-trophic behavior. MSCs expressing the fused yCD::UPRT gene convert nontoxic 5-FC to 5-fluorouracil (5-FU) on the site of tumor, thus avoiding systemic toxicity. Such cells have been designated “therapeutic mesenchymal stem/stromal cells” (Th-MSCs) (Cihova *et al*, 2011). The therapeutic potential of the MSC-driven yCD::UPRT/5-FC suicide gene therapy system was found to show a degree of effectiveness. By showing a strong bystander cytotoxic effect towards xenografts of human colorectal carcinoma cells and melanoma both *in vitro* and on nude mice *in vivo* (Kucerova *et al*, 2007; Kucerova *et al*, 2008) we demonstrated the universality of the therapeutic ability of expanded yCD::UPRT-AT-MSCs in the presence of 5-FC. Systemic administration of Th-MSCs resulted in therapeutic cells homing in on subcutaneous tumors and mediating tumor growth inhibition (reviewed in Altaner, 2008).

In a pilot preclinical study with nude mice, we demonstrated that the human yCD::UPRT-AT-MSCs were effective in significantly inhibiting subcutaneous xenografts of human bone metastatic prostate cells. The ThMSCs were administered intravenously with systemic delivery of 5-FC. Tumor regression by ThMSCs was dose dependent and repeated application improved the therapeutic outcome (Cavarreta *et al*, 2010). A positive therapeutic effect of the autologous and/or human yCD::UPRT-AT-MSCs cells was proven in the autochthonous prostate adenocarcinoma in TRAMP mice which spontaneously develop aggressive prostate cancer (Abrate *et al*, 2014). Intracranial administration of the ThMSC gene therapy system has been shown in a preclinical study to be effective in the treatment of intracerebral rat C6 glioblastoma (Altanerova *et al*, 2012; Altaner *et al*, 2013; Altaner *et al*, 2014), leading to complete tumor regression in a significant number of animals. In our preclinical studies where human melanoma cells and prostate cancer cells were implanted subcutaneously into immunocompromised nude mice, a significant tumor growth inhibition was observed when suicide gene-engineered MSCs were intravenously injected. These data were not compatible with the known bio-distribution of intravenously administered cells. Studies of bio-distribution, migration, and homing in of systemically applied MSCs revealed that 80% of cells are immediately entrapped in the lung tissue and then cleared to the liver within 1 day (Leibacher & Henschler, 2016). All these data can now be explained by the action of exosomes.

The high therapeutic potential of human yCD::UPRT-AT-MSCs can be attributed to several factors. Cumulating evidence supports the notion that soluble factors and/or exosomes secreted by yCD::UPRT-AT-MSCs exhibit a paracrine/endocrine action. In the field of regenerative medicine an increasing number of studies have shown that MSC-derived exosomes recapitulate to a large extent the immensely broad therapeutic effects previously attributed to MSCs (for a review see Phinney & Pittenger, 2017). Exosomes derived from human mesenchymal stem cells was found to promote gastric cancer cell growth, thus highlighting their tumor tropic behavior (Gu H *et al*, 2016).Tumor targeted homing of MSC exosomes enriched with miR-379 administered systemically has recently been reported. The exosomes possessed a therapeutic effect in breast cancer, mediated in part through the regulation of the cyclooxygenase-2 gene (O’Brien K P *et al*, 2018). Attention has been focused on the transfer of miRNAs by exosomes as a major mechanism of the recipient cell modifications (Neviani & Fabbri, 2015). MicroRNAs are small non-coding RNA molecules of ∼22 nucleotides and are involved in the post-transcriptional regulation of gene expression. The human genome encodes >2,500 miRNAs, which target ∼60% of mammalian genes and are abundant in a number of human cell types (Friedman *et al*, 2009).

Here we report the production of exosomes from human MSCs transduced with the yCD::UPRT and tk/*HSV* genes. The exosomes carry mRNA of the particular suicide gene in their cargo. The exosomes, upon internalization and in the presence of corresponding prodrugs, inhibited the growth of a broad range of cancer cells *in vitro*. Exosomes released from therapeutic stromal cells significantly contribute to the therapeutic efficiency of the yCD::UPRT-MSCs/5-FC and HSVtk-MSC/ganciclovir suicide gene therapy systems. With regards to quality and quantity, we did not observe any association of microRNAs originating from yCD::UPRT-MSCs/5-FC with suicide gene therapeutic effect.

## Results

A schematic overview of all steps performed in this study is presented in Figure 1. MSCs prepared from various human tissues were engineered to express the suicide genes yCD::UPRT or tk/*HSV*. CM from both suicide gene-transduced cells contain exosomes carrying mRNA of the suicide gene in their cargo. The exosomes were easily internalized by the tumor cells and in the presence of corresponding prodrugs (5-FC, ganciclovir) caused their death in a dose-dependent manner.

**Figure 1.**
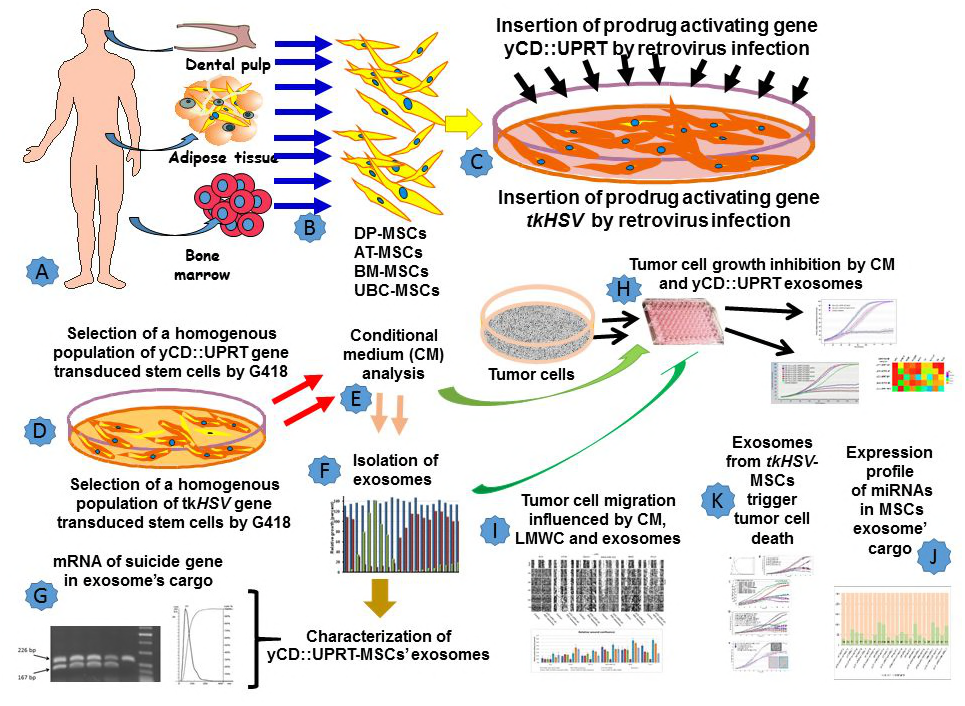
Schematic Overview of Procedures Used In This Study. **(A, B)** Isolation and expansion of MSCs from various tissues; **(C)** infection of MSCs with retrovirus carrying yCD::UPRT or tk/HSV suicide genes; **(D)** Selection of a cell population of suicide gene-transduced cells; **(E)** Harvesting of the conditional medium; **(F)** Isolation of exosomes from CM; **(G)** Detection of the mRNA of suicide genes in the exosomes’ cargo; **(H)** Tumor cell growth inhibition with CM and yCD::UPRT-exosomes; **(I)** Influence of CM, exosomes and LMWC on migration of tumor cells; **(J)** Expression profiles of miRNAs in MSCs and yCD::UPRT-MSCs exosomes; **(K)** Tumor cell growth inhibition with CM and tk/*HSV*-exosomes.

### MSCs obtained from various human tissues can be transduced with the CD::UPRT suicide gene

The design of our retrovirus vector, being a bicistronic construct with fusion yeast CD::UPRT gene separated by an internal ribosome entry site sequence from a neo gene, allows for the selection of a homogenous population of transduced cells using cell selection with the G418 antibiotic. We prepared MSCs from various human tissues and transduced them with the yCD::UPRT gene by retrovirus infection. All these genetically modified MSCs selected for their transgene presence were found to possess suicide genes integrated into their cell DNA as a DNA provirus (Fig 2 A).

**Figure 2.**
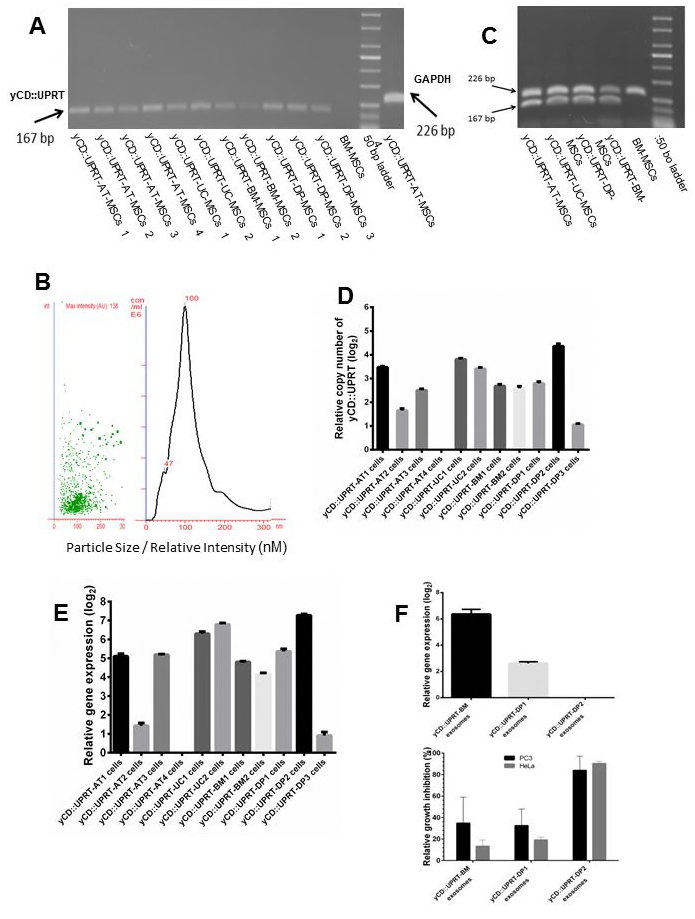
Expression of the yCD::UPRT gene in homogenous cultured yCD::UPRT-MSCs. **(A)** Isolated cellular DNA from yCD::UPRT gene transduced MSCs was PCR amplified. The presence of specific sequences was confirmed by gel resolution;**(B)** Detection of exosomes released from yCD::UPRT gene transduced MSCs by Nanosight; **(C)** Isolated DNase-treated total RNA from yCD::UPRT gene transduced MSCs was reverse transcribed and PCR amplified. The presence of specific transcripts was visualized by agarose gel electrophoresis. **(D)** qRT-PCR estimation of the relative copy number of the integrated DNA yCD::UPRT gene in cell DNA of various yCD::UPRT gene transduced MSCs. **(E)** qRT-PCR estimation of the relative expression of mRNA in exosomes from yCD::UPRT transduced MSCs; **(F)** Comparison of mRNA of the yCD::UPRT gene in the cargo of exosomes versus tumor cell growth inhibition activity.

### A homogenous population of cells transduced with yCD::UPRT gene release exosomes

It is generally observed that mammalian cells release exosomes to erradicate acquired foreign gene products or those that are highly expressed, thus maintaining cell homeostasis. We assumed that MSCs infected with the retrovirus-containing yCD::UPRT gene might release exosomes with yCD::UPRT gene products into conditional medium (CM). We prepared CM from MSCs originated from various tissues and corresponding yCD::UPRT-transduced cells (listed in Table 1) and found that all cells constantly released exosomes. Nanoparticle tracking analysis of the CM of all tested cells detected round exosomes in the range of 37 to 135 nM in diameter (Fig 2 B). The concentration of exosomes in all tested CM fluctuated at around 8.75 E9 particles/ml regardless of naive or yCD::UPRT-transduced MSCs.

**Table 1.** List of MSCs isolated from various human tissues and the corresponding yCD::UPRT gene transduced homogenous cell population used in this study

| **Human tissue type** | **Cell designation** | **Designation of yCD::UPRT gene transduced MSCs** |
| --- | --- | --- |
| Adipose tissue derived MSCs (healthy donor) | AT-MSCs (7 independent isolates) | yCD::UPRT-AT-MSCs |
| Adipose tissue derived MSCs (breast cancer patient) | AT-MSCs (breast cancer) | yCD::UPRT-AT-MSCs (breast cancer) |
| Hoffa's pad adipose tissue derived MSCs | HAT-MSCs (2 independent isolates) | yCD::UPRT-HAT-MSCs |
| Bone marrow derived MSCs (healthy donor) | BM-MSCs (3 independent isolates) | yCD::UPRT-BM-MSCs |
| Bone marrow derived MSCs (glioblastoma multiforme patient)) | BM-MSCs (GBM) (2 independent isolates) | yCD::UPRT-BM-MSCs (GBM) |
| Dental pulp derived MSCs (healthy donor) | DP-MSCs (3 independent isolates) | yCD::UPRT-DP-MSCs |
| Umbilical cord derived MSCs | UBC-MSCs (2 independent isolates) | yCD::UPRT-UBC-MSCs |
| Menstrual blood derived MSC (endometrium) | MB-MSCs | yCD::UPRT-MB-MSCs |

### The influence on tumor cell growth of conditional medium from mscs of different tissue origin compared to matched yCD::UPRT-MSCs

In order to find whether CM of yCD::UPRT-MSCs differ in any biological activity from CM of matched naive MSCs we tested how they influence growth of various human tumor cells in the presence and absence of 5-FC. The MSCs and corresponding suicide gene transduced MSCs are summarized in Table1.

Tumor cell growth was monitored in 96 microplates using a real time live cell monitoring IncuCyte system (Essen Instruments, Ann Arbor, MI), allowing for the hourly monitoring of cell growth by determining the confluence of the cells. We found that CM of naive MSCs possessed stimulating, inhibiting or no tumor cell growth activity depending on the MSC tissue origin and the types of tumor cells. The testing system for CM from yCD::UPRT-MSCs allowed us to distinguish between growth support induced by CM (5-FC is not added to CM) and tumor cell growth inhibiting activity (5-FC is added to CM). CM of yCD::UPRT-MSCs reacted similarly in the absence of 5-FC but a strong growth inhibition of tumor cells was found when 5-FC was added to CM. An efficacy comparison of the ability of CM from 5 types of yCD::UPRT-MSCs to induce tumor cell death of various tumor cell lines is presented in Figure 3E. Generally, all tumor cells that we tested, either of human or animal origin, were found to be susceptible to suicide gene therapy providing the prodrug 5-FC was added to CM.

**Figure 3.**
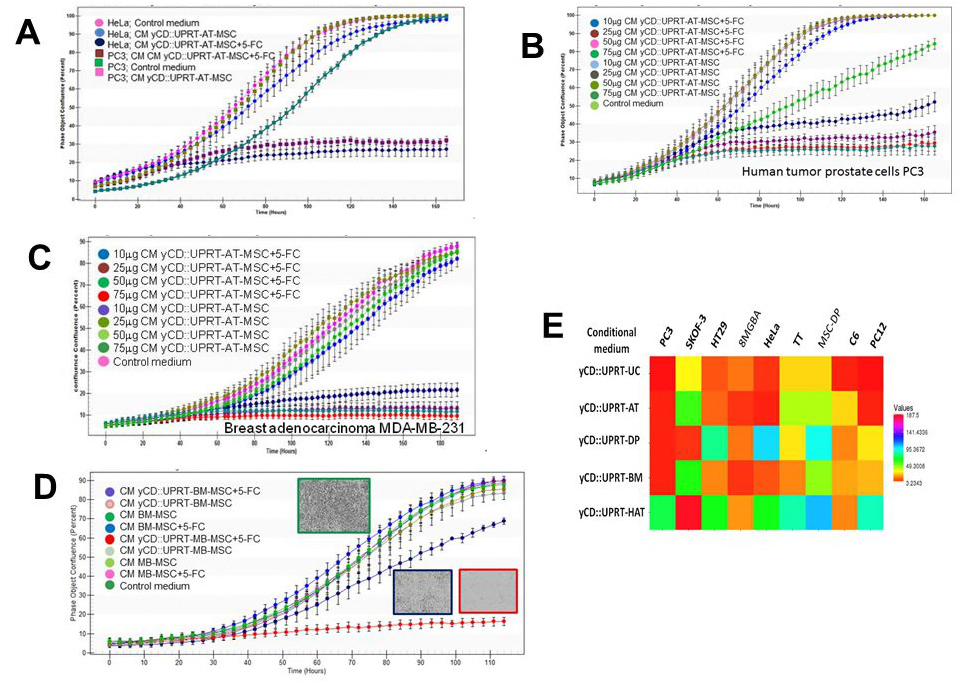
Assessment of tumor cell growth inhibition activity by exosomes from various MSCs engineered to express the yCD::UPRT gene. **(A)** Growth of PC3 cells treated with CM from yCD::UPRT-AT-MSCs in the absence of the prodrug 5-FC was stimulated, but HeLa cells were not influenced. Death was induced in both tumor cells in the presence of the prodrug; **(B)** Growth of human prostate tumor cells PC3 treated with CM from yCD::UPRT-AT-MSCs was inhibited, dose dependently, in the presence of 5-FC; **(C)** Growth of breast adenocarcinoma human tumor cells MDA-MB-231 treated with CM from yCD::UPRT-AT-MSCs was inhibited, dose dependently, in the presence of 5-FC, but in the absence of 5-FC there was no influence on their growth; **(D)** Rat glioblastoma cells C6 were effectively killed by CM from yCD::UPRT-MB-MSCs, and partially killed by yCD::UPRT-BM-MSCs. Both MSCs were isolated from the tissues of a young patient with recurrent IV grade glioblastoma; **(E)** An efficacy comparison of the ability of CM from 5 types of yCD::UPRT-MSCs to induce tumor cell death of various tumor cell lines.

Analysis of growth curves of tumor cells under the influence of CM from yCD-UPRT-transduced MSCs in the presence of 5-FC revealed that about 30 hours were needed for the mRNA of yCD::UPRT-gene to be translated intracellular into cytosine deaminase converting 5-FC to 5-FU. In the absence of 5-FC the influence of CM on the tumor cell growth could be estimated by comparing it to the control medium (Figure 3A). PC3 cells were stimulated in growth when CM from yCD::UPRT-AT-MSC was tested (Figure 3B) while MDA-MB-231 were not (Fig 3C). The tumor cell growth inhibiting effect of CM from all types of yCD-UPRT-transduced MSCs was dose dependent in all tumor cells tested (Fig 3B, Fig 3C). Comparison of tumor cell inhibiting activity of CMs prepared from yCD::UPRT-AT-MSCs isolated from healthy persons versus yCD::UPRT-AT-MSCs prepared from breast cancer patients did not reveal any differences when tested on the same tumor cells. CM from umbilical cord cells (UC-MSC) usually stimulated tumor cell growth, but the CM from corresponding yCD::UPRT-UC-MSCs were found to be highly efficient in triggering tumor cell death. The CM obtained from yCD::UPRT-MSCs inhibited the cell growth not only of human tumor cells derived from cell lines, but also of cells derived from primary human glioblastoma or rat glioma cells C6 composed mainly of glioma stem cells (Figure 3D). The exosomes of yCD::UPRT-MSCs sustained the tumor cell tropism of yCD::UPRT-MSC cells, regardless of the tissue origin of the MSCs. In agreement with our published data (Altanerova, 2017) we found that 120 minutes was a sufficient period of time for the exosome penetration and internalization into the tumor cells. The high efficiency of exosomes from yCD::UPRT-AT-MSCs in triggering the death of the human bone metastasis prostate cells PC3 is demonstrated in a video. (Supplemented file Video 1.) Real time growth inhibition of human bone metastasis prostate cells PC3 by MSC-yCD::UPRT gene exosomes in the presence and in the absence of prodrug 5-fluorocytosine. Tumour cell death induced by MSC suicide gene exosomes.

### Messenger RNA of the prodrug converting gene is present in the cargo of exosomes released from yCD-UPRT-MSC cells

We assumed that exosomes might be responsible for the tumor cell growth inhibition effect. CM from MSC-yCD::UPRT gene transduced cells was subjected to ultracentrifugation. Total RNA was isolated from pelleted exosomes, treated with DNaseI and analyzed by PCR using primers for yCD amplification. PCR analysis revealed the presence of mRNA specific to the fused y-CD::UPRT gene in all types of yCD-UPRT-MSCs analyzed (Fig 2C). Therefore, the yCD-UPRT-transduced MSCs packed the transgene mRNA into the exosome’s cargo. A similar result was obtained using an exoEasy Maxi Kit to isolate exosomes and the following exosome RNA isolation using a RNeasy Mini Kit. The internalization of exosomes into tumor cells and subsequent intracellular translation of m-RNA to an enzyme converting 5-FC to 5-FU was responsible for tumor cell death. Real time live tumor cell monitoring, treated with CM in the presence of 5-FC, revealed that the conversion takes approximately 24 – 30 hours.

### Tumor cell growth inhibition: efficacy of exosomes from yCD::UPRT gene transduced MSCs of various tissues origin

The efficiency of therapeutic exosomes derived from various types of MSCs differed in triggering tumor cell death. The yCD::UPRT-gene transduced MSCs of different tissue origin were prepared by retrovirus infection yielding cells with random integrated DNA proviruses. The presence of a neo gene linked to the yCD::UPRT gene by IRES in the retrovirus vector allowed for G418 antibiotic selection of the homogenous population of transgene containing cells. We asked whether the transgene copy number in various MSCs differed by means of qRT-PCR. We found different levels of transgene integration in cell DNA (Fig 2 D). The yCD::UPRT gene expression in the most tested cells agreed with the copy number of the integrated gene (Fig 2E). The attempts to link the level of mRNA in exosomes to the biological activity of released exosomes to cause tumor cell death by intracellular conversion of 5-FU to 5-FU in tumor cells was tested on PC3 and HeLa cells. As shown in Fig 2F the efficacy of tumor cell death induction by yCD::UPRT exosomes varied even for the same type of yCD::UPRT-MSCs and also for different isolates, as was indicated by a dental pulp MSC sample. The differences in the growth inhibition of the same exosomes were noted when tumor cells of different origin were used for testing. (Fig 2F). This suggests that either mRNA packaging in the exosome’s cargo might be cell origin related or there is variation in the penetration into tumor cells. The yCD::UPRT mRNA translation to the enzyme might be of a different efficacy in various tumor cells. Nevertheless, all yCD::UPRT gene transduced MSCs of different tissue origin, from the 27 preparations we tested, produced exosomes with the ability to induce tumor cell death in the presence of 5-FC.

### Separation of exosomes out of LMWC of the CM from yCD::UPRT-MSC cells by size-exclusion chromatography

Secretome, represented by CM from MSCs or from the corresponding yCD::UPRT gene transduced MSCs is composed of biologically active factors including a number of cytokines, chemokines, growth factors and exosomes. In order to find out whether exosomes or biological factors in the CM of yCD::UPRT gene transduced adipose tissue derived MSC (yCD-UPRT AT-MSC) possessed tumor cell growth inhibiting activity we analyzed the CM by size-exclusion chromatography on a Sepharose CL-2B column or on a Sephacryl 500 HR column. All obtained fractions were tested for their tumor killing activity in the presence/absence of 5-FC with regards to tumor cell growth activity. Human tumor prostate cells, PC3 cells, were used for testing. Figure 4A shows that the tumor cell killing activity was separated into two fractions. The majority of the tumor growth inhibiting activity was localized to the nanoparticle fractions where the highest calculated specific activity (growth inhibiting value/μg nucleoprotein) was detected. In addition, the highest number of nanoparticles was found in the same fractions by Nanosight analysis. The second fraction with low specific activity was in the lower molecular weight elution area. This tumor cell inhibiting activity could likely be contributed to by the free yCD-UPRT enzyme converting 5-FC to 5-FU or some tumor growth inhibiting factors released from MSCs.

**Figure 4.**
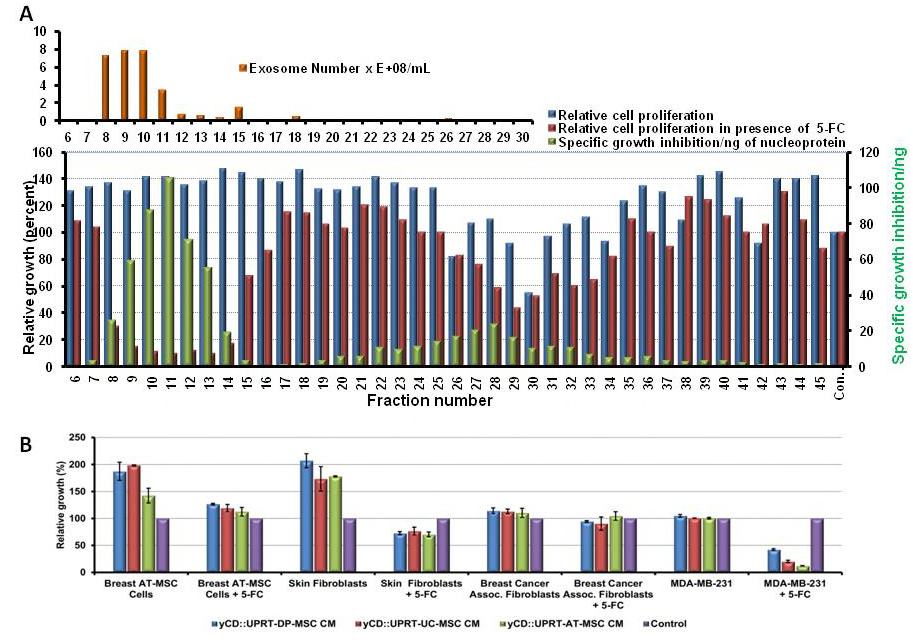
Size-exclusion Chromatography of CM from yCD::UPRT-AT-MSCs and Tumor Cell Specificity of Exosomes. **(A)** Separation of CM from yCD::UPRT-AT-MSCs; (**B**) Cell specificity of CM derived from various types of yCD::UPRT transduced MSCs.

### Tumor cell specificity of exosomes derived from different tissue-derived yCD::UPRT-AT-MSCs

In order to see whether CM from yCD::UPRT-AT-MSCs possess different efficacy of tumor cell inhibition activity we chose the following approach: we prepared explants from adipose tissue containing cancer cells of a breast cancer patient and explants from adipose tissue free of cancer cells from the same amputated mamma. Fibroblastic cells were obtained from both types of explants by cultivation over several weeks. Cells did not differ in morphology, but they differed in their ability to be induced to differentiation *in vitro*. Cells from tumor-free adipose tissue behaved as mesenchymal stem cells, while cells from adipose tissue with tumor cells were not able to differentiate into adipocytes. We designated them tumor-associated fibroblasts. In addition, fibroblasts were isolated from a human skin specimen. Comparison of relative growth of the breast cancer cells MDA-MB-231with the three types of above mentioned fibroblastic cells, all influenced by CM from adipose tissue, dental pulp and umbilical cord-derived yCD::UPRT transduced MSCs under the presence or absence of 5-FC is presented in Fig 5B. CM from all three yCD::UPRT transduced MSCs without 5-FC stimulated growth of fibroblasts from tumor-free adipose tissue and skin fibroblasts had no influence on the breast cancer associated fibroblast or breast cancer cells. A strong growth inhibition of breast cancer cells MDA-MB-231 in the presence of the prodrug was observed for all tested CMs. These data show selective internalization of breast tumor cells by therapeutic exosomes over tumor-adjacent fibroblasts, tumor associated fibroblasts and skin fibroblasts.

**Figure 5.**
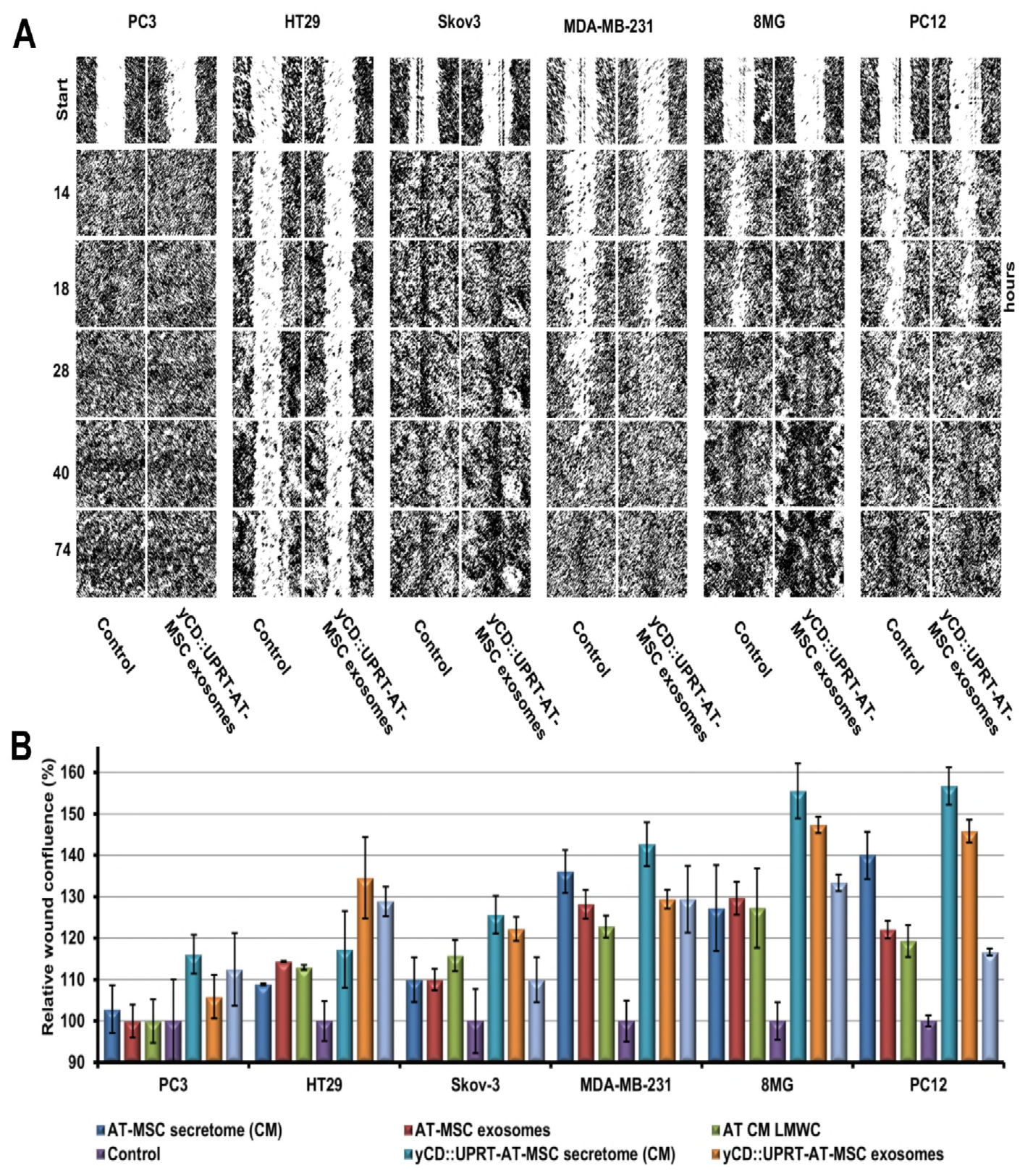
Effect of CM, Exosomes and LMWC on the Migration of Various Tumor Cells. **(A)** Effect of yCD-UPRT-AT-MSC exosomes on tumor cell wound closure; **(B)** Effect of CM, exosomes and LMWC derived from yCD::UPRT-AT-MSCs on relative wound confluence of several human tumor cells

### Effect of CM, exosomes and LMWC on the migration of various tumor cells

Separation of the exosomes from the LMWC by size exclusion chromatography allowed us to assess the effect of CM, exosomes and LWMC released from yCD::UPRT-AT-MSCs on the migration of various tumor cells. Cell migration was determined by scratch assays using a real time cell imaging system (IncuCyte live-cell ESSEN BioScience Inc, Ann Arbor, Michigan, USA), and 6 types of human tumor cells. Representative photomicrographs of tumor cell wound closure influenced by yCD::UPRT-AT-MSC exosomes in comparison with a control medium in a time (hrs) dependent fashion are presented in Figure 5A. Exosomes supported the migration of all tested tumor cells. The rate of migration was found to be dependent on the type of tumor cells. Interestingly, human colorectal carcinoma cells HT29 migrated extremely slowly while, on the other hand, human cells derived from human bone metastasis of prostate cancer were able to close the scratch in less than 14 hours.

Effects of CM, exosomes and LMWC from naive AT-MSCs and the corresponding yCD::UPRT-gene transduced cells on migration of 6 human tumor cells compared to a control medium is presented as a relative wound confluence (percent) over time (Figure 5B). Relative wound confluences of all tested cancer cells influenced by products from naivenaive AT-MSCs and yCD::UPRT-gene transduced AT-MSCs were higher compared to the influence of the control medium. The stimulation effect of CM from yCD::UPRT-gene transduced AT-MSCs was higher but not significantly different from the influence of exosomes alone. This suggests the involvement of both products in this process. The data of relative wound confluence was in good accordance with wound closure images.

### Expression of microRNAs In MSCs andcCorresponding yCD::UPRT-MSCs of various tissue origin

MicroRNAs are an important cargo of MSC exosomes. Agilent 8x60K microarray technology, able to detect 2549 human microRNAs, was used to follow the changes in miRNA expression between MSC exosomes and corresponding yCD::UPRT-MSC exosomes of various tissue origin. We observed variances in the detection rate of microRNAs between samples originating from the same donor (with and without yCD::UPRT transduct), as well as among exosomes derived from different donors and tissue types, based on the presence (Fig 6A) and expression of microRNAs (Fig. 6B). However, there was no individual microRNA specific for either MSCs or yCD::UPRT-MSC groups. Global statistical analysis focused on two questions with the following results: i) Comparison of exosomes from MSCs and corresponding yCD::UPRT-MSCs did not show any statistically significant difference regarding the change in the expression of microRNAs (BH adjusted p ‒ value < 0.05). ii) Evaluation of mean miRNA expression differences in exosomes between MSCs and yCD::UPRT-MSCs within individual types of MSCs was also negative (BH adjusted p ‒ value < 0.05). We did not detect (based on the presence as well as expression of microRNAs in exosomes) any common statistically significant difference that might suggest that deregulation of the expression of microRNAs between exosomes originating from MSCs and yCD::UPRT-MSCs was associated with therapeutic effect due to the modulation of target gene expression in tumor cells. Data of microRNA expression analysis did not show any statistically significant similarity in expression profile of exosomal microRNAs when comparing MSCs of different tissue of origin. Thus we can conclude that the expression profile of microRNAs in exosomes is unique for each tissue type.

**Figure 6.**
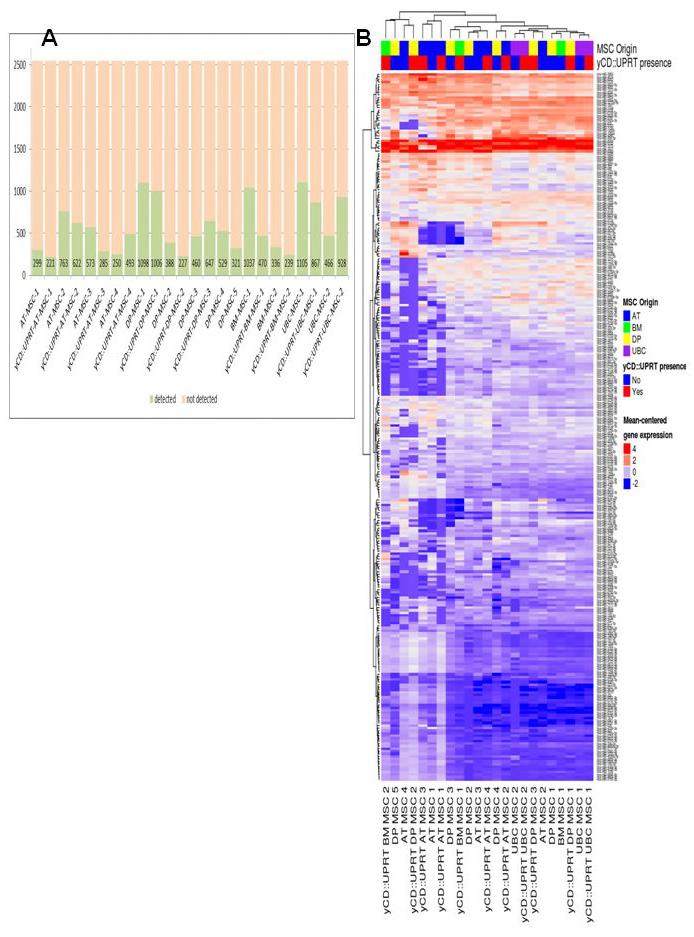
Expression profiles of differentially expressed miRNAs in MSC exosomes released from un-transduced and yCD::UPRT-gene transduced cells. **(A)** Comparison of the number of detected microRNAs (green columns) out of 2549 targeted by the microarray based on tissue types and the presence or absence of the yCD::UPRT gene. **(B)** Heat map based on the expression level of the subset (276 miRNA, expressed in at least 75% of the samples; supplementary table 1) of detected microRNAs in exosomes released from different tissue originated MSCs and corresponding yCD::UPRT gene-transduced MSCs showing no differences between the MSC and Th-MSC types (BH adjusted p - value > 0.05).

### Exosomes released from mscs expressing tk/*HSV* inhibited growth of glioblastoma cells in the presence of ganciclovir

We have shown previously that MSCs expressing thymidine kinase of the Herpes simplex virus (tk*HSV*) exert a bystander killing effect on human glioblastoma cells (Matuskova *et al*, 2010). tk *HSV* transduced adipose tissue derived MSCs (tk *HSV*-AT-MSC) released exosomes into CM. Analysis of size distribution of exosomes using NanoSight LM10 revealed the presence of nanoparticles in a range of 30 to 200 nM with a peak at 140 nM at a concentration of 7.22E9 particles/ml (Fig 7A). We tested the efficacy of the tk *HSV*-AT-MSC CM for growth support/inhibition of human glioblastoma cell line cells and primary glioblastoma multiforme cells in the absence and presence of prodrug ganciclovir. All tested glioblastoma cells were growth stimulated when CM was applied without the presence of the prodrug. When the ganciclovir prodrug was added to the CM, strong dose dependent growth inhibition was noted (Fig 7B, Fig 7C).The killing efficacy comparison of CM with corresponding tk*HSV*-AT-MSC cells on the U118 human glioblastoma cell line revealed that 10 μg of CM protein was equal to 65 cells. Similar testing using human 8MG-BA glioblastoma cells has shown higher efficiency of 10 μg of CM protein over 65 tk*HSV*-AT-MSCs (Figure 7D). Exosomes in CM from tk*HSV*-AT-MSC cells inhibited growth, not only of human glioma cells, but also rat glioblastoma cells C6. Exosomes present in CM from yCD::UPRT-AT-MSC and from tk/*HSV* cells both in the presence of the corresponding prodrugs inhibited total growth of C6 cells to the same extent. Morphological analysis of survived cells suggested a killing effect of selected glioma cell populations (Fig 7E).

**Figure 7.**
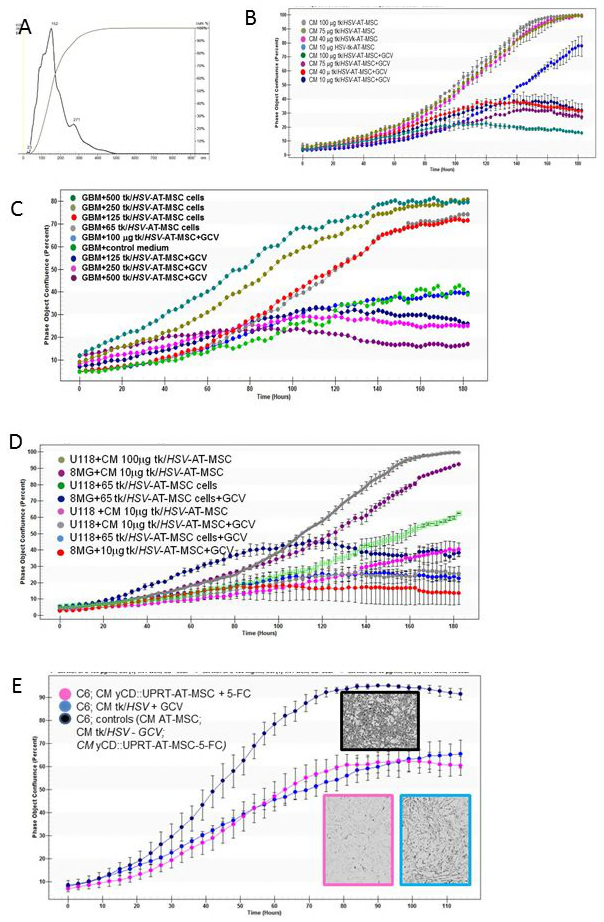
Assessment of glioblastoma cell growth inhibition activity by exosomes from AT-MSCs engineered to express thymidine kinase of the Herpes simplex virus (tk*HSV*). **(A)** Size distribution of tk/HSV-AT-MSC exosomes revealed by a NanoSight LM10 instrument; **(B)** Influence of CM from tk/HSV-AT-MSC on glioblastoma cells in the presence and absence of the prodrug ganciclovir; **(C)** CM from tk/HSV-AT-MSC dose dependent inhibited growth of human glioblastoma cells; **(D)** The CM from tk/*HSV*-AT-MSCs showed higher efficiency in growth inhibition of human 8MG-BA glioblastoma cells than tk/*HSV*-AT-MSC cells; **(E)** Growth inhibition of a selective population of rat globlastoma cells C6 treated with CM from yCD::UPRT-AT-MSC and from tk/HSV-AT-MSC cells in the presence of the corresponding prodrugs.

## Discussion

Patient survival is improving slightly for most cancer types. This progress can generally be attributed to faster diagnosis and advances in treatment. However, there is no curative cancer therapy for several solid tumors with the exception radical tumor surgery at early stages. Standard cancer therapies like radiotherapy, cytotoxic chemotherapy, even biological therapy with drugs based on monoclonal antibodies of human solid tumors and metastatic stages of the disease merely eleviate the disease symptoms and prolong the survival of cancer patients. Survival of patients with pancreatic cancer, glioblastoma, metastatic melanoma, metastatic lung tumors, high-grade serous subtype epithelial ovarian cancer, abdominal bowel and esophagus tumors treated with standard therapies have the worst outcome and the median period of survival time is extremely low. Major drawbacks of standard therapies lie in the **l**ack of tumor specificity of anti-cancer drugs, therapy not being addressed to tumor-initiating stem cells and in the emergence of drug-resistant cell subpopulations after chemotherapy. Wilhelm at al., 2016 reviewed the drug delivery literature from the past decade and estimated the anti-cancer drug delivery efficiency to tumors. Only 0.7% (as a median) of an injected dose of drug in a form of nanoparticles ends up in the tumor.

The natural behavior of mesenchymal stem cells in targeting tumors led us to develop prodrug gene therapy mediated by MSCs more than ten years ago (Kucerova *et al*, 2007). In our preclinical studies with human melanoma cells (Kucerova et al., 2010), prostate cancer cells (Cavarreta et al2010) implanted subcutaneously into immune-compromised nude mice along with intravenously injected ThSc displayed significant tumor growth inhibition. We interpreted the high tumor inhibiting activity as bystander effect of 5-FU released from injected therapeutic cells. These data are not compatible with the currently known bio-distribution measurements of intravenously injected labeled MSCs. The cells are immediately entrapped in lung tissue and then cleared to the liver within one day (Leibacher & Henschler, 2016).

Here we report evidence that MSCs with the yCD::UPRT gene integrated in the cell DNA under strong retrovirus promoter express mRNA of the transgene and pack it into the cargo of released exosomes. We have proved that all yCD::UPRT gene transduced human MSCs derived from various tissues like adipose, bone marrow, dental pulp, umbilical cord and menstrual blood derived endometrial regenerative cells release exosomes containing mRNA of the suicide gene in the exosome’s cargo. Thus the efficacy of prodrug gene therapy for cancer mediated by the yCD::UPRT-MSCs/5-FC system and by the tk*HSV*-MSC/ganciclovir system act not only through a bystander effect but mainly through exosomes that internalize tumor cells. It was reported that intrinsic properties of tumor cells, like expression of the ABC transporters and the ability of gap-junctional intercellular communication (GJIC), have a key impact on the bystander effect mediated by genetically engineered mesenchymal stem cells. The cytotoxic effect mediated by the yCD::UPRT-MSCs/5-FC system and by the tk*HSV*-MSC/ganciclovir system was evaluated in direct co-culture experiments. The efficacy of both therapeutic systems was assigned by the authors to GJIC, the expression of enzymes involved in drug metabolism and proteins responsible for drug transport (Matuskova *et al*, 2012). The authors did not consider the existence of MSC-exosomes with suicide gene mRNA; therefore, the explanation only concerned the bystander effect. The involvement of therapeutic MSC exosomes with paracrine/endocrine activity causing intracellular tumor cell death has to be taken into consideration. These exosomes could easily communicate with cancer cells through specific receptor-ligand interactions. They internalize tumor cells bringing not only mRNA of the suicide gene, but other mRNA and regulatory microRNAs as well, being able to induce and/or modify various molecular paths in the cells (Validi *et al*, 2007). Combined action of the bystander effect and therapeutic exosome internalization into tumor cells will take place when cells are involved in the therapy of metastases.

MSC-exosomes with suicide genes represent a novel anti-cancer drug, targeted at the tumor, internalizing tumor cells – a therapy with intracellular activity. We have called them MSC therapeutic exosomes (MSCThE). The discovery of MSCThE with paracrine/endocrine activity is an important step toward cell-free prodrug gene therapy. This finding explains our previous studies where we have shown a curative therapy of rat glioblastoma treated with intracerebral administration of human yCDy-UPRT-AT-MSCs cells (Altanerova *et al*, 2012; Altaner *et al*, 2013; Altaner *et al*, 2014). Accumulating evidence indicates that cancer therapy using therapeutic MSC exosomes has multiple advantages over cell therapy. Similarly, an advantage of exosomes over cells in clinical trials using exosomes for patients suffering from steroid-refractory acute graft-versus-host disease (acute GvHD) or chronic kidney disease with improvement reports was comprehensively discussed by Giebel *et al*, 2017. So far, MSC-exosome administration appears to be safe in humans. CM or exosomes are stable after intravenous administration and exhibit a superior safety profile. Since MSCs have the remarkable tendency to home in on tumors, exosomes produced by MSCs may retain the homing properties of their parent cells. Dental pulp derived MSCs being of neural crest-derived cells might serve as an example. We have recently shown that dental pulp-derived MSCs can migrate to intracerebral glioblastomas after intranasal administration (Altanerova et al, 2016).

MicroRNAs as a cargo of exosomes were considered as a major mechanism of the recipient cell modifications (Neviani & Fabbri, 2015). We have been dealing with the question of whether the antitumor effect of exosomes derived from yCD-UPRT-MSCs can be fully attributed to the presence of mRNA originating from the inserted construct, or whether miRNA molecules may play a role in antitumor effects. Therefore, we compared the spectrum of miRNAs present in the exosomes derived from a yCD-UPRT-MSC-culture medium with the miRNA spectrum in exosomes derived from MSCs without a construct. Based on the presence as well as expression of microRNAs in exosomes, we have not found any common statistically significant difference between exosomes originating from MSCs and yCD-UPRT-MSCs. Therefore we do not think that exosomal miRNAs could be associated with therapeutic effect by the modulation of target gene expression in tumor cells. This does not exclude the possibility of the active involvement of exosomal miRNAs in the gene regulation in recipient tumor cells (O’Brien *et al*, 2017).

A number of studies in the field of regenerative medicine has shown that nanoparticles produced by MSCs exert their therapeutic effects in several diseases, suggesting that MSC-derived exosomes are responsible for therapeutic effect. MSC-derived nano vesicles may confer a stem cell-like phenotype to injured cells with the consequent activation of endogenous self-regenerative programmers. Regenerative medicine mediated by MSC cells is changing to cell-free therapy mediated by MSC-derived exosomes (Phinney & Pittenger, 2017). Exosomes represent carriers with virus-like properties that can efficiently regulate gene expression in recipient cells. They differ from viruses in not being infectious.

The composition of exosomes released from MSCs might be purposely modified to create therapeutically interesting exosomes for cancer treatment (Balyasnikova *et al*, 2010). Recently we reported the dual action of MSCs-yCD-UPRT-MSC exosomes possessing, besides mRNA of the suicide gene, iron oxide nanoparticles as well. These nanoparticles were able to act both as a prodrug dependent therapeutic and hyperthermia inducing factor when tumor cells were exposed to an alternating magnetic field (Altanerova U *et al*, 2017). Pascucci *et al*, 2014 reported a new approach for drug delivery proving that if MSCs are treated with Paclitaxel, the drug is incorporated into exosomes. Therapeutically active exosomes released from yCD-UPRT-MSCs supports the earlier observation that MSC driven prodrug gene therapy might be the modality for treatment of diffuse intrinsic pontine glioma, the most malignant and dismal of cancers in children (Choi *et al*, 2012; Roger *et al*, 2010).

Collectively, we believe that for an innovative cancer therapy to be curative, it must differ from standard therapies which merely alleviate the symptoms of the disease. Any novel cancer therapy has to be targeted not only at the tumor, but also specifically at tumor cells and the therapy has to act intracellularly. This could avoid the side effects of standard therapies. The therapeutic exosomes possessing the mRNA message of suicide genes and/or classical chemotherapeutic drugs in their cargo might be the novel approach to curative cancer therapy. In numerous preclinical *in vivo* studies with MSCs engineered to express suicide genes applied intravenously we have come to understand (Cavarretta *et al*, 2010; Altanerova *et al*, 2012; Altaner *et al*, 2014) that the exosomes released from these cells were clearly behind the promising positive outcomes. Nevertheless, new animal studies will be performed with therapeutic exosomes only administered intravenously. These could give the decisive answer on the usefulness of prodrug gene therapy for cancer mediated by MSC exosomes.

## Materials and Methods

### Culture isolation and maintenance of MSCs

All donors of adipose tissue, bone marrow, blood platelets, and other tissue specimens used for isolation and propagation of MSCs were informed about the nature of the study and provided their written informed consent. We have previously described all MSC isolation procedures. AT-MSCs were isolated by collagenase digestion (Kucerova *et al*, 2007). BM-MSCs, MB-MSCs were isolated using density gradient centrifugation (Percoll separating solution, density 1.077g/ml) (Altaner *et al*, 2013), DP-MSCs and UBC-MSCs were isolated from tissue fragments adhered to plastic tissue culture dishes (Stanko *et al*, 2014). For the expansion of all kinds of MSCs, the cells were seeded at 4000 cells/cm^2^ in plastic dishes (Corning Life Sciences) and grown with medium exchange every 2-3 days. Adherent cells were split after reaching confluence with 0.05% trypsin/EDTA (Gibco). All MSCs used for the experiments were up to fifth passage or less. MSC cultures were grown in a complete culture medium DMEM low glucose (1 g/L) supplemented with 5% human platelet extract (PE), antibiotic antimycotic solution (Gibco Life Technologies) and incubated at 37 °C in a humidified atmosphere with 5% CO_2_. All tumor cell lines were maintained in DMEM high glucose (4 g/L) supplemented with 5% of fetal bovine serum.

### Preparation of yCD::UPRT-retrovirus-producing cells

Recombinant retrovirus possessing the yCD:;UPRT fused gene was prepared as described previously (Kucerova L, *et al*, 2007). Briefly, GP+E-86 helper cells were transfected with plasmid designated pST2 containing bicistronic retrovirus construct with the yCD:;UPRT gene separated by IRES sequence from the neo gene. Virus-containing medium from G418-resistant ecotropic helper cells was used to infect amphotropic GP+envAM12 helper cells. Three to five ping-pong rounds of repeated infections were performed to obtain cells producing recombinant retrovirus particles with mixed envelope glycoproteins as described (Hlavaty *et al*, 1999). The cells were cloned to obtain a highly virus producing clone for further use. The virus-containing medium for transduction was collected from a selected high virus-producing cell clone of semi-confluent cultures of GP+envAM12/pST2 cells. The virus containing medium without serum (24 hour harvest) was filtered through 0.45-μm filters, and used either fresh or kept frozen at minus 80°C until use.

### Cell transduction with retrovirus

All types of MSCs transduced with the yCD::UPRT-gene by retrovirus infection was performed as previously described (Kucerova *et al*, 2007). Briefly, subconfluent MSC cultures were transduced three times in three consecutive days with a virus-containing medium from GP+envAM12/ pST2 cells supplemented with 100 μg/ml protamine sulfate. Cells were cultivated in selection media containing a pretested concentration of G418 able to kill naive cells (over a range of 0.4 to 1.2mg/ml) of G418 for several days and expanded to obtain the same batch of cells for the experiments conducted *in vitro*.

The following tumor cell lines were used: human colon adenocarcinoma cells HT-29; breast adenocarcinoma cells MDA-MB-231; ovarian carcinoma cell line SKOV-3, medullary thyroid carcinoma TT, bone metastasis prostate cells PC3; cervical cancer cells HeLa; glioblastoma cells 8MG-BA, malignant glioma cell U118; primary glioblastoma multiforme cells; human fibroblast cells, rat pheochromocytoma cells PC12 and rat glioblastoma cells C6.

### Cell growth assessment using a IncuCyte live cell monitoring system

Cell growth was monitored by “real time *in vitro* micro-imaging” using the IncuCyte system (EssenInstruments, Ann Arbor, MI). The IncuCyte system allows for the hourly monitoring of cell growth by determining the confluence of the cells and displaying the morphologic changes associated with treatment. Most cell viability data obtained from the Incucyte system were in good accordance with data obtained by subjecting the plates to the CellTiter 96 Aqueous One Solution Cell Proliferation Assay (Promega).

### Isolation of exosomes

Conditioned medium was the source of exosomes. CM was collected from 90% confluent cell cultures with 90 -97% cell viability assessed by Trypan-Blue exclusion (Sigma–Aldrich, St. Louis, MO). The cells were washed with PBS to remove any debris and cultured over 24–48 hrs. in the growth supplement deficient medium. Conditioned media collected from the culture dishes were centrifuged at 800g for 5 min to remove cells and cell debris. CM harvested from all gene transduced AT-MSCs was collected on ice and filtered through a 0.22 μm syringe filter.

Exosomes were separated from the CM by ultracentrifugation at 110,000g for 15 h in a fixed angle 70Ti rotor (Beckman Coulter, Pasadena, CA, USA) to pellet exosomes. The exosomal pellet was washed once in PBS and then resuspended in 200 μl of phosphate-buffered saline (PBS; Life Technologies) and stored at -80° C. All centrifugation steps were performed at 4° C.

### PCR and qRT-PCR analysis

Total DNAs were extracted from all types of yCD::UPRT gene transduced MSCs and from BM-MSCs (as negative control) using a GeneJET RNA Purification Kit (Thermo Scientific). Standard PCR was performed in 10 μl 5x Colorless GoTaq Reaction buffer (Promega), 1 μl PCR Nucleotide mix (10 mM each), 2 μl of each primer for CD::UPRT (10 μM), 0,25 μl GoTaq DNA polymerase (5U/ μl, Promega), 6 μl DNAs (7 ng/μl) and 28,75 μl nuclease-free water. Thermal cycling conditions for 35 cycles were: initial denaturation at 95°C for 2 minutes, denaturation at 95°C for 45 sec, annealing at 50 °C for 30 sec, extension at 72°C for 30 sec, final extension at 72° C for 5 min.

As positive control of amplification, PCR with primers for GAPDH was performed with the same DNA, using the same thermal cycling conditions, except the annealing temperature was 53°C for GAPDH primers.

The following primers were used:

5´-ATGGACATTGCCTATGAGGA-3´ (forward) and

5´-TTCTCCAGGGTGCTGATCTC-3´ (reverse) for yCD (167 bp),

5´-GAAGGTGAAGGTCGGAGTC-3´(forward) and

5´-GAAGATGGTGATGGGATTTC-3´ (reverse) for GAPDH (226 bp).

PCR products were visualized by agarose electrophoresis (2%) and identified using a GeneRuler 50 bp DNA Ladder (Thermo Fisher Scientific).

Total RNAs were isolated from all types of yCD::UPRT gene tranduced MSCs and from BM-MSCs (as negative control) using a GeneJET RNA Purification Kit (Thermo Scientific). RNAs were treated with Ambion RNase-free DNase I (Invitrogen). Reverse transcriptase reactions were performed using a Maxima First Strand cDNA Synthesis Kit for RT-qPCR (Thermo Scientific). The presence of CD::UPRT in cDNA was confirmed by PCR with the same conditions as for DNA.

Real-time PCRs for DNA as well as for cDNA were carried out with a Quant Studio 3 Real-Time PCR System (Applied Biosystems) using a TaqMan gene expression system with TaqMan Universal Master Mix II, according to the manufacturer’s instructions. Results were analyzed by QuantStudio^TM^ Design & Analysis Software. Gene expression was compared using delta cycle threshold (ΔCt = Ct_CD::UPRT_-Ct_RNASE_ P) values for the respective cells examined where RNASE P expression was taken as an endogenous reference. Analysis was performed in triplicates and data expressed as mean ± SD. yCD::UPRT-AT-MSCs were taken as a reference sample. Change in gene expression was calculated according to the formula: *RQ* = 2^‒ΔΔ*Ct*^. Differences between cell line expression were tested using the one-way ANOVA and Tukey’s multiple comparisons test. P-values <0.05 were considered to indicate statistically significant differences. The data were presented as log_2_ of relative quantity (RQ) of analysed DNA or cDNA.

### Tumor cell migration assay

Tumor cells were grown to confluence in 96-well culture plates (Essen BioSCIENCE, Ann Arbor, Michigan,USA) for 24 h. Cell migration was assessed by scratch assays, made using a 96-pin WoundMaker. The wells were washed with PBS to remove any debris and incubated in the presence of CM, exosomes and LMWC. Cell migration was visualized using a real-time cell imaging system (ESSENBioScience Inc, Ann Arbor, Michigan, USA). Cells were imaged every 2 hours to monitor the treatment-induced cell migration.

### Isolation of RNA from released exosomes in conditional medium

Isolation of exosomes and subsequent isolation of total RNA was achieved by two different methods. The first method of isolation of exosomes was performed using an exoEasy Maxi Kit (Qiagen, Hilden, Germany) from 1 ml of conditioned media (CM) and subsequent isolation of RNA from exosomes was done with an exoRNeasy Serum / Plasma Starter Kit (Qiagen, Hilden, Germany) according to the company’s instructions. The second was the isolation of exosomes using a kit from Exiqon (Vedbaek, Denmark) miRCURY™ Exosome Isolation based on precipitation of exosomes from the 1 ml of CM, and isolation of total RNA by a miRCURY RNA Isolation Kit according to the company’s instructions. To determine individual miRNA expression the Qiagen kit was selected due to higher yields of RNA.

### Detection of microRNAs

Agilent SurePrint G3 Unrestricted miRNA 8x60k microarray (G4872A, design ID 070156) was used for detection of 2549 human miRNAs based on the miRBase database release 21.0. In total, 24 CM samples divided into 8 groups were compared for differential expression of miRNAs – AT-MSCs (4 CM samples), yCD::UPRT-AT-MSCs (4), BM-MSCs (2), yCD::UPRT-BM-MSCs (2), DP-MSCs (5), yCD::UPRT-DP-MSCs (3), UBC-MSCs (2) and yCD::UPRT-UBC-MSCs(2).

RNA isolated from the conditioned media was labeled and hybridized to the slides following the manufacturer’s protocol (miRNA Microarray System with miRNA Complete Labeling, Version 3.1.1) without the optional steps. Hybridized slides were scanned immediately using a SureScan Microarray Scanner (Agilent) with automatic settings. Scanned slides were analyzed by Feature Extraction Software (Agilent) and resulting signal intensities were further processed in R software. After quality control and data normalization, expression data were tested using a random effect model, moderated t-statistic and simple Bayesian model to assess differential expression of individual miRNAs. After adjustment based on 10% Benjamini and Hochberg’s false discovery rate, miRNAs with adjusted P-values < 0.05 were considered to be differentially expressed.

## Acknowledgments

This study was supported by a grant awarded to CA by the Slovak League against Cancer and by the National Sustainability Program I (NPU I) Nr. LO1503 provided by the Ministry of Education Youth and Sports of the Czech Republic. We thank Dusan Guba MD, Institute of Medical Cosmetics, Bratislava, for providing us with material for AT-MSC isolation.

## Author contributions

CA, and UA designed and implemented the study; UA, JJ, and KB isolated and expanded cells, realized cell growth inhibition analyses; PP and VR designed and implemented molecular analyses; CA realized cell transduction experiments, MZ performed Nanosight analyses; PP, MP, and AT contributed the work on microRNA; CA, UA, MP, AT, and JK interpreted data; CA, UA, and MP wrote the manuscript; All authors contributed to and have approved the final manuscript.

## Abbreviations

MSCs: mesenchymal stem cells
DP-MSCs: dental pulp MSCs
AT-MSCs: adipose tissue MSCs
BM-MSCs: bone marrow MSCs
UBC-MSCs: umbilical cord MSCs
yCD::UPRT: yeast cytosine deaminase::uracil phosphoribosyl transferase suicide fusion gene
tk/HSV: thymidine kinase of the Herpes Simplex Virus.
CM: conditioned medium
LMWC: low molecular weight components
mRNA: messenger RNA
miRNA: micro RNA.

